# Integrate and generate single-cell proteomics from transcriptomics with cross-attention

**DOI:** 10.1101/2025.01.28.635217

**Authors:** Jiankang Xiong, Shuqiao Zheng, Fuzhou Gong, Liang Ma, Lin Wan

## Abstract

**Motivation:** Single-cell RNA sequencing (scRNA-seq) and cellular indexing of transcriptomes and epitopes by sequencing (CITE-seq) have experienced rapid advancements in recent years, accompanied by the development of numerous methods for analyzing scRNA-seq and CITE-seq data. These innovations have enabled deeper insights into cellular heterogeneity and functional phenotypes. However, analyzing scRNA-seq and CITE-seq data within a unified framework remains a significant challenge in the field of single-cell analysis. Specifically, this challenge centers on two primary objectives: aligning scRNA-seq and CITE-seq cells within an integrated representation space and generating antibody-derived tag (ADT) measurements for scRNA-seq cells.

**Results:** By incorporating interrelationships between cells into a deep generative model with cross-attention, we introduced scProca to integrate and generate single-cell proteomics from transcriptomics. scProca delivers state-of-the-art performance in both integration and generation tasks across benchmark datasets. Furthermore, scProca can accommodate cells across experimental batches, showcasing its flexibility in complex experimental contexts.

**Availability:** The code of scProca is available at https://github.com/xiongbiolab/scProca, and replication for this study is available at https://github.com/ZzzsHuqiaAao/scProca-reproducibility.

## 1 Introduction

The rapid progression of single-cell RNA sequencing (scRNA-seq) technologies has revolutionized molecular biology by enabling the comprehensive evaluation of transcriptome profiles at an unprecedented resolution (Svensson *et al*., 2018). Nonetheless, scRNA-seq only captures a single layer of the intricate regulatory machinery that governs cellular function and signaling (Heumos *et al*., 2023). Consequently, there is a growing interest in the development of single-cell multiomic techniques to enhance our understanding of cellular identity by providing multiple perspectives on molecular states (Efremova and Teichmann, 2020). By combining highly multiplexed protein marker detection with unbiased transcriptome profiling, cellular indexing of transcriptomes and epitopes by sequencing (CITE-seq) leverages oligonucleotide-conjugated antibodies to simultaneously quantify counts of RNA expression and surface protein abundance in single cells via the sequencing of antibody-derived tags (ADTs), offering a unique opportunity to bridge the transcriptome with functional protein information (Todorovic, 2017; Lischetti *et al*., 2023), and ultimately enabling a more comprehensive characterization of cellular phenotypes beyond transcriptome measurements alone (Stoeckius *et al*., 2017).

CITE-seq enhances the identification of specific protein markers associated with particular transcriptomics profiles, improving the accuracy and precision of cellular classification (Frangieh *et al*., 2021; Rebuffet *et al*., 2024). This enables researchers to gain deeper insights into cellular functions and states (Liu *et al*., 2023), while foster a more nuanced understanding of cellular heterogeneity, allowing for clearer distinctions between closely related cell populations (Lischetti *et al*., 2023). These advancements have, in turn, driven the development of unsupervised analysis methods specifically designed for CITE-seq data (Athaya *et al*., 2023). Recently, new approaches have emerged to leverage the strengths of both technologies to create a more comprehensive understanding of cellular functions (Hu *et al*., 2024). Specifically, these methods focuses on two primary objectives: aligning the scRNA-seq and CITE-seq cells within an integrated representation space, and generating ADT measurements for scRNA-seq cells.

For the alignment of scRNA-seq and CITE-seq cells within an integrated representation space, the deep generative model totalVI probabilistically models CITE-seq data as a composite of biological and technical factors, including protein background and batch effects, extending the analysis of CITE-seq to the integration of scRNA-seq and CITE-seq (Gayoso *et al*., 2021). Alignment can also be accomplished through transfer learning. For instance, scArches employs transfer learning and parameter optimization to facilitate efficient, decentralized, iterative reference building and contextualization of new datasets with existing references (Lotfollahi *et al*., 2022). More generally, alignment can be categorized under multi-batch, multi-modal mosaic integration. In this context, scVAEIT introduces a probabilistic variational autoencoder model for mosaic integration, which merges data sources containing both shared features across datasets and features unique to a single data source (Du *et al*., 2022). UINMF presents a nonnegative matrix factorization approach that incorporates a metagene matrix to handle both shared and unique features in single-cell data integration (Kriebel and Welch, 2022). scMoMaT integrates single-cell multi-omics data in the mosaic integration scenario using matrix tri-factorization (Zhang *et al*., 2023). StabMap enhances the stability of single-cell mosaic data integration by inferring a data topology based on shared features and projecting cells onto reference coordinates (Ghazanfar *et al*., 2024). totalVI and UINMF have been shown to excel beyond other methods in aligning scRNA-seq and CITE-seq cells within an integrated representation space, as demonstrated in the latest benchmarking by Hu *et al*. (2024).

Among the methods, only totalVI, scArches, and scVAEIT can simultaneously generate, impute, or predict ADT measurements from RNA measurements. Additionally, cTp-net proposes a transfer learning framework based on deep neural networks (Zhou *et al*., 2020), Guanlab-dengkw uses kernel ridge regression (Lance *et al*., 2022), and sciPENN develops a single-cell imputation protein embedding neural network (Lakkis *et al*., 2022), all of which are capable of generation. Furthermore, the classic toolkit Seurat can also transfer data from reference to query using a transfer learning framework (Stuart *et al*., 2019). totalVI and scArches have been shown to outperform other methods in predicting protein abundance, as demonstrated in the latest benchmarking by Hu *et al*. (2024).

By incorporating interrelationships between cells into a deep generative model, we introduced scProca to integrate and generate single-cell proteomics from transcriptomics. With cross-attention, scProca effectively addresses challenges of posterior inference caused by variability in input types, where some cells are RNA-seq and others are CITE-seq. Extensive benchmarking demonstrated that scProca achieved state-of-the-art performance in both integration and generation tasks. Furthermore, scProca can accommodate cells across experimental batches, showcasing its flexibility in complex experimental contexts.

## 2 Methods and Materials

### 2.1 scProca

We introduce scProca, a deep generative model to integrate and generate **s**ingle-**c**ell **pro**teomics from transcriptomics with **c**ross-**a**ttention (Fig.**1**A). scProca incorporates a variational auto-encoder (VAE) enhanced with cross-attention, enabling the inference of batch-corrected, integrated latent variables from scRNA-seq and CITE-seq data. These latent variables can then be used in the optimized generative process to generate the expression of ADT that were not sequenced in the scRNA-seq data.

**Figure 1.**
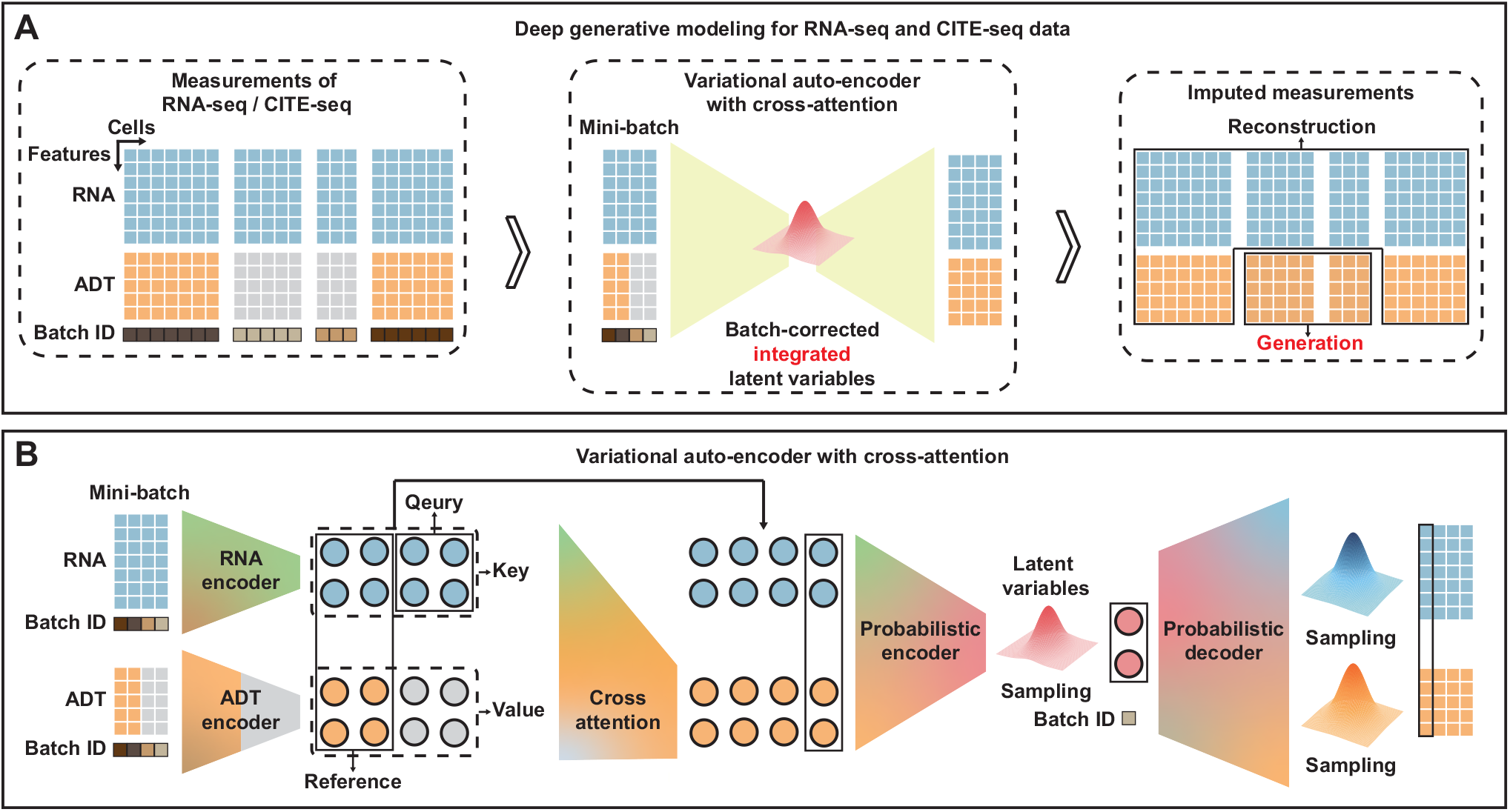
(A) Schematic representation of scProca within the framework of deep generative models. scProca is capable of inferring batch-corrected, integrated latent variables from scRNA-seq and CITE-seq data, and generating the expression profiles of ADT for scRNA-seq cells. (B) The variational auto-encoder with cross-attention introduced in scProca. Cross-attention is used to incorporate CITE-seq cells as references, completing representation of scRNA-seq cells in the ADT embedding space.

The VAE enhanced with cross-attention, effectively performs posterior inference in scenarios where input types vary, with some cells being scRNA-seq and others being CITE-seq (Fig.**1**B). For both omics, specialized encoders are employed to extract useful information from raw features into their respective embedding spaces. When a cell is scRNA-seq, lacking ADT measurements, it bypasses the ADT encoder. Instead, cross-attention is used to incorporate CITE-seq cells as references, completing its representation in the ADT embedding space. Subsequently, the embeddings from both omics are passed through a probabilistic encoder to infer the posterior distribution of the integrated latent variables. To mitigate batch effects, an adversarial loss is applied to the latent variables.

When applied to large-scale datasets, the computation of cross-attention becomes challenging. To address this, we employ a stochastic optimization strategy, leveraging the commonly used mini-batch trick in neural network training.

### 2.2 Deep generative process for scProca

Let 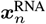 and 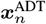 denote the expression profiles (column vector) of the *n*th cell for transcriptomics and proteomics respectively. These expression variables are assumed to be generated through the following process:

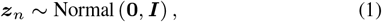

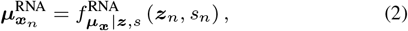

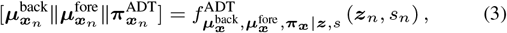

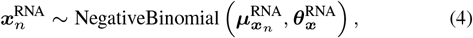

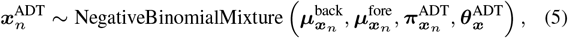

where *s*_*n*_ is the batch annotation for the *n*th cell. Equation (1) defines the latent variable ***z***_*n*_, representing the intrinsic cellular variability of the *n*th cell and is sampled from a standard normal distribution. Equations (2)-(5) describe the conditional distributions of the transcriptomics and proteomics expression profiles. Following Gayoso *et al*. (2021), the expression profiles 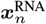 and 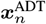 are modeled a Negative Binomial distribution and a Negative Binomial Mixture, respectively, with the learnable inverse dispersion parameters ***θ***_***x***_. The mean parameters 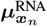 for the Negative Binomial distribution are obtained via a neural network 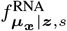. Similarly, the background means 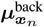, foreground means 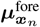, and mixing rates parameters 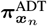 for the Negative Binomial Mixture are obtained via another neural network 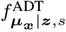. These networks take as input the latent variable ***z***_*n*_ and the batch annotation *s*_*n*_. For clarity, their learnable parameters of the neural networks 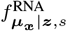 and 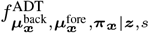 are omitted in the notation.

### 2.3 Posterior inference for the latent variables

Given the generative model for the transcriptomics and proteomics expression profiles, we aim to infer the posterior distribution of the latent variables ***Z*** = [***z***_1_, ⊆, ***z***_*N*_] conditioned on the observed data 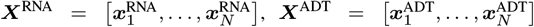 and the batch annotations ***s*** = [*s*_1_, ⊆, *s*_*N*_]^*T*^. The exact posterior *p* (***Z***| ***X***^RNA^, ***X***^ADT^, ***s)*** is intractable due to the integral of *p* (***X***^RNA^, ***X***^ADT^ |***s*)**. To approximate this posterior, we use a variational distribution *q* ( ***Z***|***X***^RNA^, ***X***^ADT^, ***s*)** and employ the evidence lower bound (ELBO):

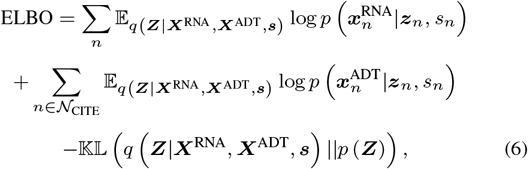

where *𝒩*_CITE_ ⊆ [1, ⊆, *N*] is the subset of cells being CITE-seq. The prior distribution 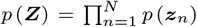 is given by Equation (1). The conditional generative distribution 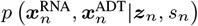 factorizes as:

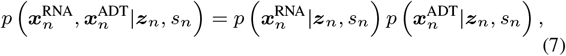

with each term modeled by Equations (4) and (5), respectively.

The variational posterior distribution *q* ( ***Z***|***X***^RNA^, ***X***^ADT^, ***s*)** is expressed as the product of independent terms for each data point:

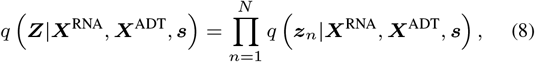

where each *q* (***z***_*n*_|***X***^RNA^, ***X***^ADT^, ***s*)** is chosen to be a Normal distribution with a diagonal covariance matrix:

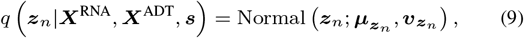

where Normal 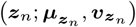 represents the probability of ***z***_*n*_ under the Normal distribution with mean 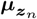 and diagonal covariance 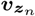.

The first two terms of the evidence lower bound (ELBO) correspond to the negative reconstruction loss, in the context of autoencoders. The last term represents the Kullback-Leibler (KL) divergence regularization on the variational posterior distribution.

To account for inter-cellular interactions and address the lack of proteomics measurements in scRNA-seq cells, we propose an encoding process with cross-attention (Vaswani, 2017), to infer the mean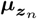 and standard deviation 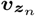 of the variational posterior distribution. This is achieved through the following encoding steps:

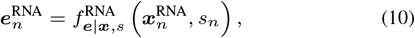

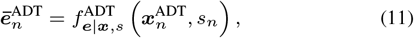

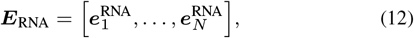

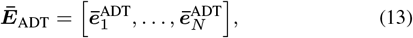

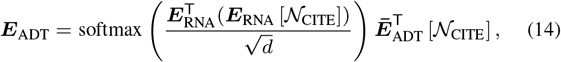

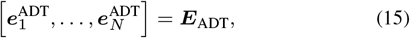

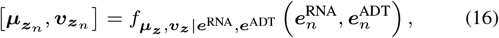

where *d* denotes the dimensionality of the latent space. In Equations (10) and (11), the functions 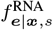 and 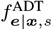 are neural networks that encode the RNA and ADT data for each cell respectively, conditioned on the batch annotation *s*_*n*_, to obtain the omics-specific embedding of the *n*th cell. As proteomics measurements may be unavailable for scRNA-seq cells, we use a cross-attention operation to impute missing proteomics representation in the embedding space, as described in in Equation (14), where ***E***_RNA_ [*𝒩*_CITE_] denotes the selection of the rows in ***E***_RNA_ indexed by *𝒩*_CITE_. Finally, the embeddings 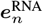 and 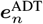 are integrated by a probabilistic neural network 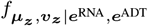 to produce the parameters of the variational posterior distribution 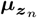 and 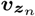.

### 2.4 Adversarial learning

To align the latent representations of cells across different experimental batches, we introduce a batch-adversarial loss. This loss encourages the model to remove batch-specific information from the latent variable ***z***_*n*_, as well as embeddings 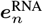 and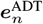. We accomplish this by training discriminators to predict the batch origin of the latent variable and embeddings, using neural network classifiers 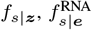and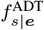, where 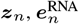 and 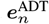 are input to the respective discriminators. The latent variable and embeddings are trained to fool these discriminators. When the discriminators and the variational posterior distribution reach equilibrium, the discriminators cannot distinguish the batch origins of the latent variable and embeddings, effectively mitigating batch effects.

The adversarial loss is formulated as:

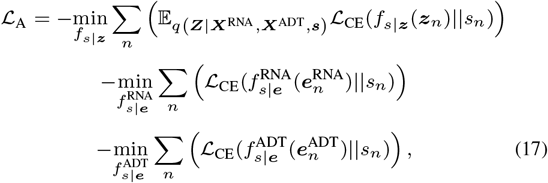

where *ℒ* _CE_ denotes the cross-entropy loss between the predicted and true batch labels. The discriminators aim to minimize the cross-entropy loss, while the variational posterior aims to maximize it to fool the discriminators.

### 2.5 Reconstruction from embeddings

To improve the interpretability of the learned embeddings, we impose an auxiliary reconstruction loss on both the RNA and ADT embeddings, 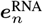 and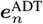, ensuring that they can reconstruct the observed measurements. The reconstruction loss is defined as:

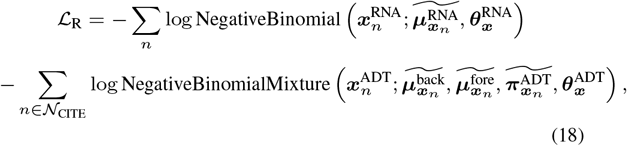

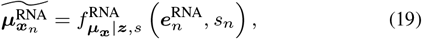

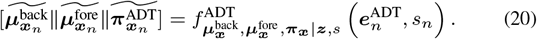

### 2.6 Training of scProca

The overall loss *ℒ* for scProca is based on the ELBO loss *ℒ* _ELBO_ = −ELBO, the reconstruction loss on the embeddings *L*_R_, and the batch-adversarial loss *ℒ* _A_, formulated as:

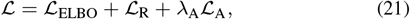

where *λ*_A_ is the coefficient for the batch-adversarial loss, with its default value set to 30.

Given the presence of adversarial loss, adversarial iterative optimization is employed. The learnable parameters are divided into two sets: one set, denoted as ***G***, includes the parameters of the assumed prior conditional distribution and the designed variational posterior distribution, i.e.,

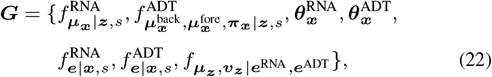

and the other set, denoted as ***D***, includes the parameters of the discriminators, i.e.,

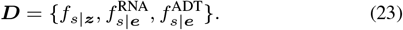

When ***G*** is fixed, the dual loss *L*_D_ of *L*_A_, is minimized with respect to ***D*** as follows:

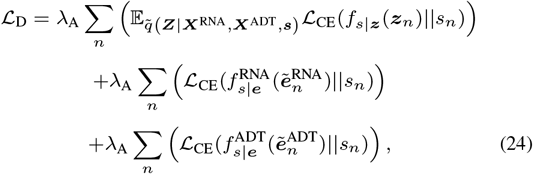

where 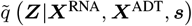, 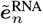 and 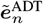 represent the fixed ***G*** used for optimizing ***D***. When ***D*** is fixed, the overall loss *ℒ* is minimized with respect to ***G*** as follows:

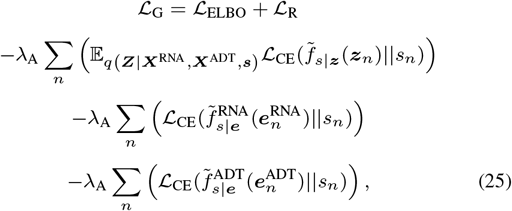

where 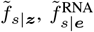 and 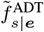 represent the fixed ***D*** used for optimizing ***G***.

The expectation terms in both losses *ℒ* _G_ and *ℒ* _D_ are approximated using a single sample. Both losses are minimized using the Adam optimizer with an initial learning rate of 4e-3 and a weight decay of 1e-6.

### 2.7 Integration of multi-batch transcriptomics and proteomics

Due to the absence of proteomics measurements for scRNA-seq cells, integrating multi-batch cells into a common space presents a significant challenge. To address this, we utilize batch-adversarial learning, where the latent variable ***z***_*n*_ helps mitigating batch effects. Additionally, since the latent variable *z*_*n*_ encodes the intrinsic cellular variability of individual cells, we use the estimation of the integrated latent variable, 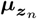, inferred from the variational posterior distribution, as a low-dimensional representation of the *n*th cell. This representation can be subsequently used for further analyses, such as assessing batch effects and evaluating clustering accuracy.

### 2.8 Generation of missing proteomics

In accordance with the generative model assumptions, cellular expression profiles are generated from the latent variable, regardless of whether proteomics measurements are available. Thus, we estimate the latent variable, 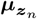, from the variational posterior distribution and use this to generate the proteomics measurements for the *n*th scRNA-seq cell as 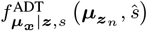, where *ŝ* denotes the annotation of the anchor batch to which the proteomics measurements are generated.

### 2.9 Stochastic optimization and inference

To efficiently handle large-scale datasets, we employ the mini-batch trick, which is used by both scProca and other neural network-based methods. At each iteration, a random mini-batch of data (512 samples) is sampled, eliminating the need to process the entire dataset at once and thus reducing memory requirements. The iterative objective functions from Equations (24) and (25) are continuous and end-to-end differentiable, enabling the use of automatic differentiation to perform stochastic optimization. To mitigate the variability introduced by random division of mini-batches, posterior inference is repeated multiple times (100 times) to obtain an averaged estimation.

## 3 Results

### 3.1 scProca achieves state-of-the-art performance in integration on benchmark datasets

We assessed the performance of scProca in comparison with other competitive integration methods across four benchmark datasets, following the evaluation metrics introduced in recent benchmarking studies by Hu *et al*. (2024) (Supplementary Note 1 and Supplementary Note 2). The visualization of low-dimensional representations shows that, compared to other integration methods, scProca performs better on the DOGMA dataset by more effectively mixing scRNA-seq and CITE-seq cells, while also clustering different cell types together (Fig.**2**A). In more comprehensive quantitative metrics (Supplementary Fig.S1), scProca achieves state-of-the-art performance, outperforming other methods in terms of both biological variation conservation and batch effect removal scores (Fig.**2**B), as well as the overall integration score (Fig.**2**C). Among the other methods, UINMF and totalVI rank second and third, respectively, consistent with the benchmark results from Hu *et al*. (2024). UINMF shows acceptable performance in batch effect removal, it lags behind in biological variation conservation, ranking at the bottom, while totalVI performs moderately, with both biological variation conservation and batch effect removal scores near the median of all methods. scProca also achieves state-of-the-art integration performance on the REAP, SLN, and BMMC dataset (Supplementary Fig.S2).

**Figure 2.**
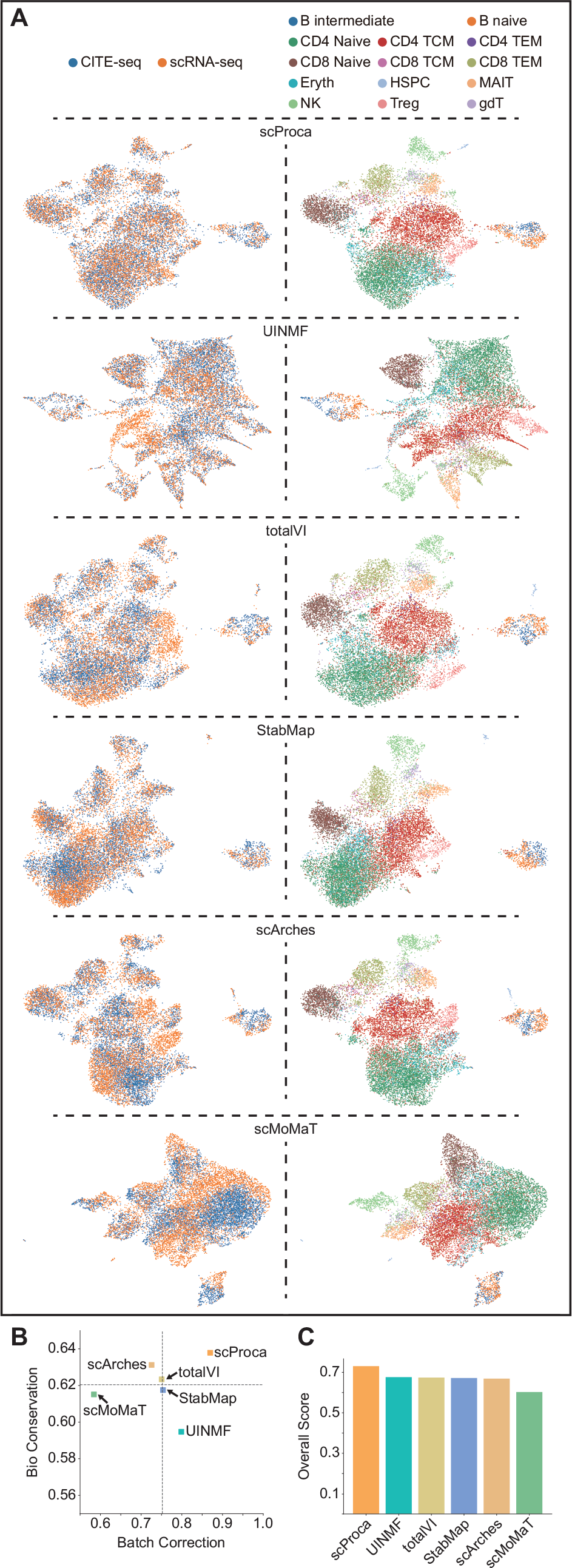
(A) UMAP visualization of the low-dimensional representations obtained by scProca and five other integration methods on the DOGMA dataset, labeled by batch annotations and cell types. (B)The scatter plot of biological variation conservation and batch effect removal scores for integration. The dashed lines represent the median scores of all methods. (C) Overall integration scores across all methods.

### 3.2 scProca achieves state-of-the-art performance in generation on benchmark datasets

We also assessed the performance of scProca in comparison with competitive generation, imputation or prediction methods on the benchmark datasets, also following the benchmark metrics introduced by Hu *et al*. (2024) (Supplementary Note 2). Scatter plots of the two main generation metrics, PCCs and CMDs, show that scProca achieves state-of-the-art performance on the DOGMA dataset. Along with totalVI, it is one of the only two methods where both dimensionality metrics are above the median value for all methods (Fig.**3**A and Fig.**3**B). The ranking index indicates that scProca and totalVI tie for first place in overall performance, followed by scArches (Fig.**3**C), which is also consistent with the benchmark results from Hu *et al*. (2024). When broken down by individual metrics, sciPENN ranks highly for the RMSE metric (Fig.**3**D), but performs poorly in others, especially for cell-cell PCC and CMD metrics (Fig.**3**E and Fig.**3**F). Similarly, Seurat shows poor performance in both the cell-cell PCC and CMD metrics, despite excelling in protein-protein PCC metric (Fig.**3**G). Although Seurat performs well on the protein-protein PCC metric, it also struggles with protein-protein CMD metric (Fig.**3**H). Guan-Dengkw, which performs poorly on RMSE metric, excels in protein-protein CMD metric. These results suggest that most methods tend to focus on specific aspects of performance, often at the expense of others. In contrast, both scProca and totalVI provide more balanced performance across all metrics. scProca also achieves state-of-the-art generation performance on the REAP (Supplementary Fig.S3), SLN (Supplementary Fig.S4), and BMMC (Supplementary Fig.S5) dataset.

**Figure 3.**
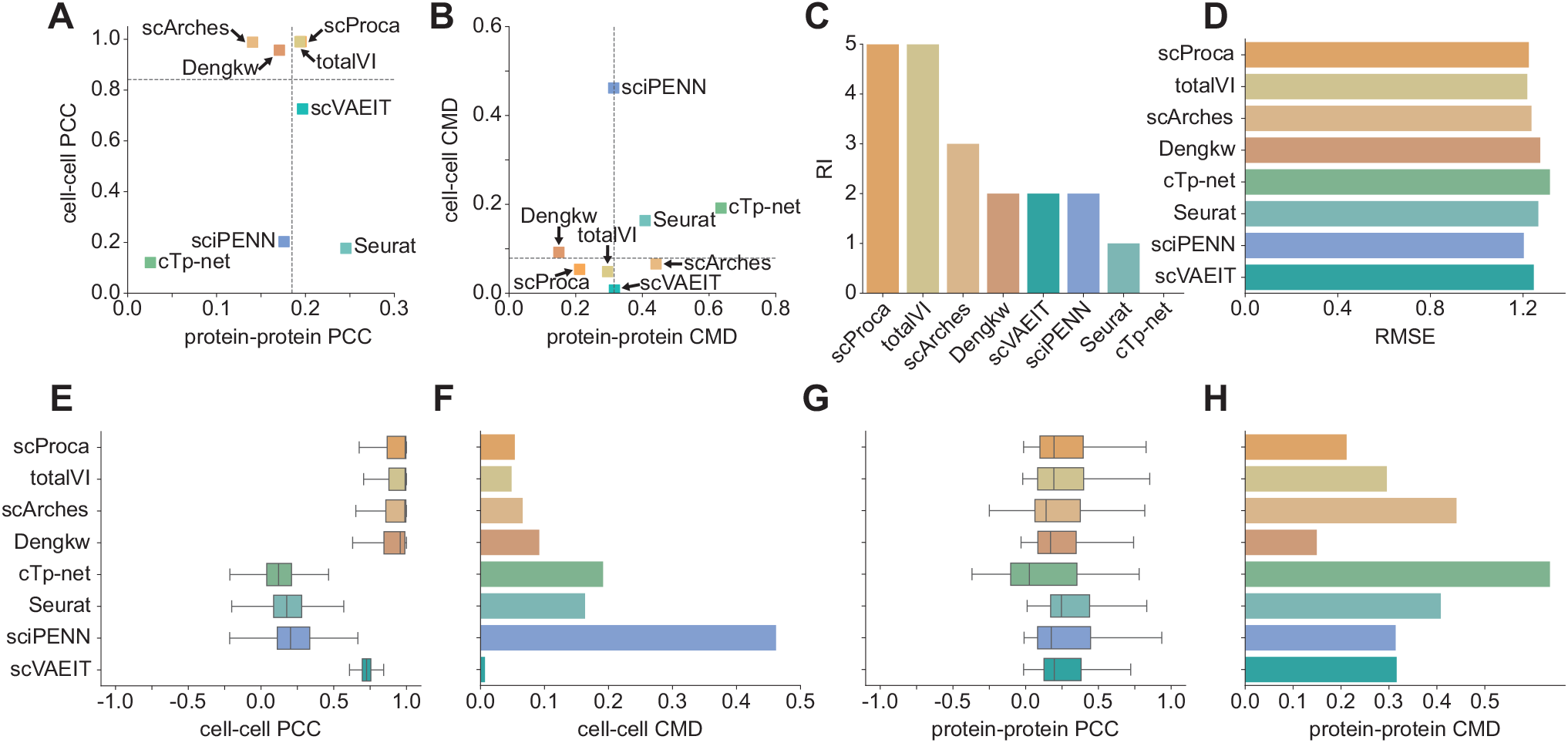
Scatter plots showing (A) cell-cell PCC versus protein-protein PCC and (B) cell-cell CMD versus protein-protein CMD between the ADT measurements of CITE-seq cells and the predicted ADT measurements of scRNA-seq cells obtained by scProca and seven other generation, imputation, or prediction methods on the DOGMA dataset. The dashed lines are the median values of all methods. (C) Overall RI scores across all methods. (D) RMSE, (E) cell-cell PCC, (F) protein-protein PCC, (G) cell-cell CMD, and (H) protein-protein CMD values across all methods.

### 3.3 scProca accommodates cells across experimental batches

In previous benchmarking, analysis is limited to reference and query batches derived from distinct datasets. However, experimental batches may also originate from the same dataset. For example, the SLN Dataset in the benchmark is divided into two datasets, but each dataset can be further subdivided into two distinct experimental batches.

To extend this analysis, we investigated scProcas performance in scenarios where only one experimental batch SLN111-D1 from SLN dataset contained proteomics expression profiles, while the proteomics profiles of the other three experimental batches were omitted, leaving them as scRNA-seq datasets. We found that scProca not only mitigated batch effects across different datasets (Fig.**4**A), but also in this scenario, effectively integrated data when the refined experimental batches were treated as sub-batches. Despite only one experimental batch serving as CITE-seq data, scProca was also able to harmonize the data (Fig.**4**B). Moreover, even with a reduced number of effective CITE-seq cells, scProca maintained its ability to cluster different cell types effectively (Fig.**4**C), without compromising its capacity to conserve biological variation (Fig.**4**D). Detailed structures of layouts revealed that, without adequately dividing the data into experimental sub-batches, the integration of latent variables across datasets appeared mixed on a dataset-wide scale but led to the separation of the same cell type based on experimental batch. In contrast, when the data was organized into sub-batches, scProca effectively grouped cells of the same type across different batches, even when only one batch served as the CITE-seq reference. This conclusion is further supported by the ARI and NMI metrics (Supplementary Fig.S6A). Additionally, in generating ADT measurements, scProca demonstrated consistent performance and did not degrade even with fewer reference cells (Supplementary Fig.S6B). This experiment allowed for a more comprehensive evaluation of scProcas ability to integrate transcriptomics and proteomics measurements across different experimental contexts and its capacity to generate proteomics data for scRNA-seq cells under more complex conditions.

**Figure 4.**
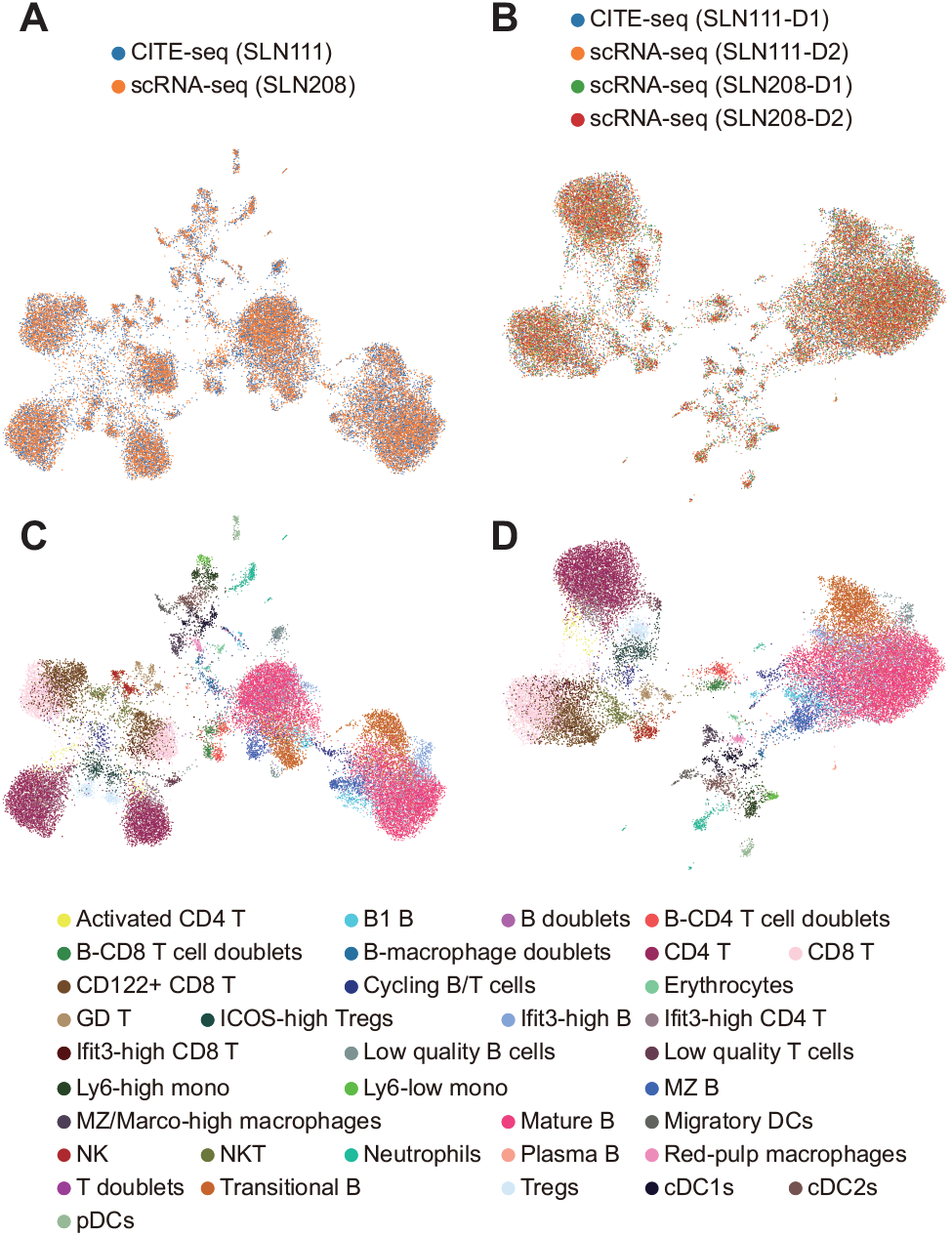
UMAP visualization of the low-dimensional representations obtained by scProca, colored by batch annotation and cell types. Color by batch annotation: (A) Only SLN111-D1 serves as CITE-seq data, while the others serve as scRNA-seq data. (B) Both SLN111-D1 and SLN111-D2 serve as CITE-seq data, and both SLN208-D1 and SLN208-D2 serve as scRNA-seq data. Color by cell types: (C) Same setup as (A). (D) Same setup as (B).

## 4 Discussion and Conclusion

In this study, we introduced scProca, a novel deep generative model, to integrate and generate single-cell proteomics from transcriptomics data. By using cross-attention, scProca effectively addresses key challenges in jointly posterior inferring of integrated latent variable for both scRNA-seq and CITE-seq cells. Our results demonstrated that scProca achieved state-of-the-art performance in both integration and generation tasks across multiple benchmark datasets consisting of scRNA-seq and CITE-seq cells. Furthermore, scProca can accommodate cells across experimental batches, showcasing its flexibility in complex experimental contexts.

In recent years, alongside the development of single-cell transcriptomics and proteomics, other single-cell omics have also been refined with advances in sequencing technologies, enabling researchers to gain a more comprehensive understanding of single cells. One of the cutting-edge fields in single-cell research is the integration and generation of more general multi-batch, multi-omics mosaic data. scProca offers strong scalability, allowing additional omics information to be incorporated in a manner similar to proteomics, enabling more comprehensive variational posterior inference. This is an area we plan to explore further in our future work. Spatial omics has also emerged as a highly popular research field. scProca can be extended to model spatial CITE-seq and spatial transcriptomics data by incorporating spatial information or imposing spatial constraints during posterior inference. This allows a natural extension into spatial omics research, which is another key direction for our future studies.

