## Supplementary data for "Integrate and generate single-cell proteomics from transcriptomics with cross-attention"

### Supplementary Information

Jiankang Xiong[1,3,†] Shuqiao Zheng[1,3,†] Fuzhou Gong[1,3] Liang Ma[2,3,*] Lin Wan[1,3,*]

[1]NCMIS, Academy of Mathematics and Systems Science, Chinese Academy of Sciences, Beijing 100190, China

[2]Institute of Zoology, Chinese Academy of Sciences, Beijing 100101, China

[3]University of Chinese Academy of Sciences, Beijing 100049, China

[*]To whom correspondence should be addressed. (Liang Ma) and (Lin Wan). [†]The authors wish it to be known that, in their opinion, the first two authors should be regarded as Joint First Authors.

### Contents

**Supplementary Methods and Materials** **3**

Supplementary Note 1: Benchmark datasets . . . . . . . . . . . . . . . . . . . . . . . 3

Supplementary Note 2: Benchmark metrics . . . . . . . . . . . . . . . . . . . . . . . 4

**Supplementary References** **8**

**Supplementary Figures** **11**

### Supplementary Methods and Materials

#### Supplementary Note 1: Benchmark datasets

All four datasets are from the recent benchmarking study by Hu et al. (2024). Among all the data provided, we selected those used for predicting protein abundance in the inter-dataset scenario. Each dataset contains sub-datasets from the same organ/tissue. In total, there are five eligible datasets, but one of them does not provide cell type information. Since we need cell type information to validate whether integration can conserve biological variation, we excluded this dataset.

The remaining four datasets include two human PBMC datasets, which were generated using different sequencing technologies. To distinguish between them, we named them according to their respective technologies: the REAP dataset and the DOGMA dataset. The other two datasets are from the spleen and lymph nodes, referred to as the SLN dataset, and from BMMC, referred to as the BMMC dataset.

In the context of the benchmarking study, Datasets 1 (Hao et al., 2021)-2 (Luecken et al., 2021) correspond to the BMMC dataset, Datasets 21-22 (Gayoso et al., 2021) correspond to the SLN dataset, Datasets 23-24 (Peterson et al., 2017) correspond to the REAP dataset, and Datasets 25-26 (Mimitou et al., 2021) correspond to the DOGMA dataset. For detailed information on each dataset, please refer to the benchmarking study.

Following the protocol outlined in the benchmarking study, the entire benchmark is conducted under the reference/query framework. In the BMMC data, Dataset 2 is used as the reference dataset, which retains both RNA and ADT measurements with a total of 90,261 cells. Dataset 1 is used as the query dataset, which only contains RNA measurements with 33,454 cells. In the SLN data, Dataset 22 serves as the reference dataset with 15,820 cells, and Dataset 21 serves as the query dataset with 16,828 cells. In the REAP data, Dataset 24 serves as the reference dataset with 4,330 cells, and Dataset 23 serves as the query dataset with 3,152 cells. In the DOGMA data, Dataset 26 serves as the reference dataset with 6,139 cells, and Dataset 25 serves as the query dataset with 7,624 cells.

#### Supplementary Note 2: Benchmark metrics

1. The Pearson Correlation Coefficient (PCC) is defined by Equation (S1):

$$\mathrm{PCC}(\mathbf{x}, \mathbf{y}) = \frac{\sum_{i=1}^{n}(x_i - \bar{x})(y_i - \bar{y})}{\sqrt{\sum_{i=1}^{n}(x_i - \bar{x})^2}\sqrt{\sum_{i=1}^{n}(y_i - \bar{y})^2}} \tag{S1}$$

When calculating cell-cell PCC, $x_i$ and $y_i$ represent the abundance of proteins in cells $x$ and $y$, respectively. $\bar{x}$ and $\bar{y}$ represent the average values of $\{x_i\}$ and $\{y_i\}$, respectively. For the calculation of protein-protein PCC, $x_i$ and $y_i$ are the protein abundances of cell $i$ for the proteins $x$ and $y$, respectively.

2. CMD is commonly used to measure the difference between two correlation matrices, $\mathbf{R}_1$ and $\mathbf{R}_2$, and a lower CMD value indicates a better result (Hu et al., 2021; Huang et al., 2018; Wang et al., 2019). The CMD is given by Equation (S2):

$$d(\mathbf{R}_1, \mathbf{R}_2) = 1 - \frac{\mathrm{trace}(\mathbf{R}_1\mathbf{R}_2)}{\|\mathbf{R}_1\|_F\|\mathbf{R}_2\|_F}, \tag{S2}$$

where $\mathrm{trace}(\mathbf{R}_1\mathbf{R}_2)$ represents the trace of matrix $\mathbf{R}_1 \times \mathbf{R}_2$ and $\|\cdot\|_F$ is the Frobenius norm of a matrix. In this study, each element in the correlation matrices $\mathbf{R}$ is the PCC value between two cells or two proteins.

3. The RMSE quantifies the difference between the predicted values ($\mathbf{X}$) and the true values ($\hat{\mathbf{X}}$)(Gayoso et al., 2021; Lakkis et al., 2022). To ensure comparability, both the predicted and true values are normalized using the same method and rescaled using $z$ scores. The RMSE is mathematically defined as:

$$\mathrm{RMSE} = \|\mathbf{X} - \hat{\mathbf{X}}\|_F, \tag{S3}$$

where $\|\cdot\|_F$ is the Frobenius norm of a matrix.

4. To assess the concordance between known cell-type labels and the cell clusters identified by the K-Means algorithm, we use the Adjusted Rand Index (ARI) (Hubert and Arabie, 1985) and Normalized Mutual Information (NMI) (Strehl and Ghosh, 2002) as our primary metrics. Given the two clustering results $U$ and $V$, following four quantities are calculated: $a$) number of objects in a pair placed in the same group in $U$ and in the same group in $V$; $b$) number of objects in a pair placed in the same

group in $U$ and in different groups in $V$; $c$) number of objects in a pair placed in the same group in $V$ and in different groups in $U$; and $d$) number of objects in a pair placed in different groups in $U$ and in different groups in $V$. ARI has been proposed in the form of:

$$\mathrm{ARI} = \frac{\binom{n}{2}(a+d) - [(a+b)(a+c)+(c+d)(b+d)]}{\binom{n}{2} - [(a+b)(a+c)+(c+d)(b+d)]}. \tag{S4}$$

NMI is defined as $I(U,V)/\max(H(U),H(V))$, where $I(U,V)$ is the mutual information between $U$ and $V$, and $H(U), H(V)$ represent the entropy of the clustering results $U$ and $V$, respectively.

$$I(U,V) = \sum_{p=1}^{P}\sum_{q=1}^{Q} \frac{U_p \cap V_q}{N} \log \frac{N|U_p \cap V_q|}{|U_p| \times |V_q|}, \tag{S5}$$

where $|U_p|$ and $|V_q|$ denote the cardinality of the $p$th cluster in $U$ and the $q$th cluster in $V$, respectively. The entropy of each cluster assignment is calculated as follows:

$$H(U) = -\sum_{p=1}^{P} \frac{|U_p|}{N} \log \frac{|U_p|}{N}, \tag{S6}$$

$$H(V) = -\sum_{q=1}^{Q} \frac{|V_q|}{N} \log \frac{|V_q|}{N}. \tag{S7}$$

5. The average silhouette width (ASW) (Rousseeuw, 1987) metric is used to gauge the precision of cell-cell distances calculated by each integration algorithm. ASW is an indicator of how well a cell matches with others in its cluster (intra-cluster similarity) and how distinct it is from cells in the closest different cluster (inter-cluster dissimilarity). The silhouette width for each cell is computed using the formula:

$$\frac{b-a}{\max(a,b)},$$

where $a$ represents the average intra-cluster distance, and $b$ represents the average nearest-cluster distance. Averaging all the silhouette widths of a set of cells yields the ASW, which ranges between -1 and 1. In our analysis, we leverage ASW scores based on cell-type labels (cASW) and batch labels (bASW) to evaluate each algorithm's effectiveness in conserving biological variation and removing batch effects, respectively. A higher cASW value signifies improved accuracy in cell-type separation, whereas a lower bASW indicates more effective correction of batch effects. To ensure a standardized evaluation, we transform the cASW and bASW values using linear transformations (in line with the method used by Luecken et al. (2022)) so that higher values consistently indicate superior performance for both the cASW and bASW.

6. The LISI (Korsunsky et al., 2019) is used to assess the results of our integration algorithms in terms of cell-type separation (denoted as cLISI) and batch mixing (referred to as iLISI). In the context of our study, a lower cLISI value is indicative of more effective cell-type separation, signifying enhanced conservation of biological variation. Conversely, a higher iLISI value reflects better integration of different batches, pointing to more successful removal of batch effects. To maintain consistency in our evaluation criteria and ensure that higher values of both iLISI and cLISI represent improved performance, we apply linear transformations to these values (Luecken et al., 2022).

7. The KNN graph connectivity (GC) (Büttner et al., 2019) metric is used to evaluate the connectivity between cells within each cell type in a KNN graph. This metric is given by the equation(S8):

$$\mathrm{GC} = \frac{1}{N}\sum_{i=1}^{N}\frac{\max(m_i)}{n_i}, \tag{S8}$$

where $N$ is the total number of cell types, $n_i$ is the cell number of cell type $i$, and $\max(m_i)$ is the cell number of the largest connected subgroup of cell type $i$ in the KNN graph.

8. PCR (Büttner et al., 2019) is used to measure the BER for multi-omics integration algorithms. The PCR is defined according to Equation (S9):

$$\mathrm{PCR} = \sum_{i=1}^{n}\mathrm{var}(A|\mathrm{PC}_i)\cdot R^2(\mathrm{PC}_i|\mathrm{batch}), \tag{S9}$$

where $A$ can be the RNA expression matrix, the chromatin accessibility matrix, the protein abundance matrix, or the low-dimensional embedding matrix generated by an integration algorithm. $\mathrm{PC}_i$ is the $i$-th principal component of $A$, $\mathrm{var}(A|\mathrm{PC}_i)$ is the variance of $A$ on $\mathrm{PC}_i$, and $R^2(\mathrm{PC}_i|\mathrm{batch})$ signifies the squared correlation coefficient between $\mathrm{PC}_i$ and the batch labels of cells.

9. We use the kBET (Büttner et al., 2019) as a metric to quantify the extent of BER by each integration algorithm. kBET assesses the similarity between two key components: the composition of batch labels among the $k$-nearest-neighbors of a cell ($C_{\mathrm{KNN}}$) and the overall batch labels composition across all cells ($C_{\mathrm{batch}}$). Ideally, in a scenario where the batch effect has been effectively eliminated, $C_{\mathrm{KNN}}$ should be equal to $C_{\mathrm{batch}}$, resulting in a kBET value of 1. We calculate the kBET value for each algorithm's integration results using scIB (Luecken et al., 2022) with default settings.

10. The isolated label score (ILS) (Luecken et al., 2022) is used to assess the effectiveness of mosaic integration algorithms in embedding cell connectivity graphs into a low-dimensional space and isolating specific cell types that are present in only a subset of data batches. Specifically, for any given cell type $i$ that occurs in $k_i$ batches, the ILS is determined by averaging the ASW values for cell types that are present in $k_{\mathrm{min}}$ batches. Here, $k_{\mathrm{min}}$ represents the smallest number among all $k_i$ values.

11. Ranking index (RI) is used to gauge the overall performance of each algorithm. The RI value of algorithm $i$ is defined according to Equation (S10):

$$\mathrm{RI}_i = \sum_j B(\nu_{ij}), \tag{S10}$$

where $B$ is the Heaviside function, that is,

$$B(x) = \begin{cases} 0, & x < 0 \\ 1, & x \geq 0, \end{cases}$$

and $B(\nu_{ij})$ represents whether algorithm $i$ is one of the top-performing algorithms when using metric $j$ for comparison. For metrics where a lower value signifies better performance, such as CMD and RMSE, we define $\nu_{ij} = \mathrm{median}(Y_{\cdot,j}) - Y_{ij}$. In contrast, for metrics where a higher value signifies better performance, such as PCC and AUROC, we define $\nu_{ij} = Y_{ij} - \mathrm{median}(Y_{\cdot,j})$. Here, $Y_{ij}$ refers to the value of the $j$-th metric for the $i$-th algorithm, and $Y_{\cdot,j}$ represents the array of values for the

$j$-th metric across all algorithms being evaluated. The dataset-specific rank index quantifies an algorithm's relative performance within a specific dataset, based on the number of metrics for which it ranks in the top 50%. This method is a nuanced variation of the RI, which is computed across all datasets. For example, if the algorithm totalVI ranks among the top 50% for three specific metrics (such as cell-cell PCC, protein-protein PCC, RMSE) in a dataset, its dataset-specific rank index for that dataset would be assigned as 3.

12. The BVC score is a key metric for evaluating the performance of each integration algorithm on a given dataset. It is calculated as the mean of several metrics: ARI, NMI, cASW, cLISI, and ILS. The BVC score is instrumental in evaluating how well an algorithm preserves biological variation across datasets, and these metrics are normalized using the scaling method outlined by scIB (Luecken et al., 2022).

13. The BER score for each integration algorithm on a given dataset is the average of five key metrics: kBET, GC, bASW, iLISI, and PCR. The BER score assesses an algorithm's ability to effectively mitigate batch effects. Similarly to the BVC score, the values of kBET, GC, bASW, iLISI, and PCR are rescaled for BER score calculation, following the transformation methodology of scIB (Luecken et al., 2022).

14. The overall score for each integration algorithm on a given dataset is the weighted average of the BVC and BER score (Luecken et al., 2022), following the equation:

$$\text{Overall} = 0.6 \times \text{BVC} + 0.4 \times \text{BER}. \tag{S11}$$

Malte D Luecken, Maren Büttner, Kridsadakorn Chaichoompu, Anna Danese, Marta Interlandi, Michaela F Müller, Daniel C Strobl, Luke Zappia, Martin Dugas, Maria Colomé-

Tatché, et al. Benchmarking atlas-level data integration in single-cell genomics. *Nature Methods*, 19(1):41–50, 2022.

Eleni P Mimitou, Caleb A Lareau, Kelvin Y Chen, Andre L Zorzetto-Fernandes, Yuhan Hao, Yusuke Takeshima, Wendy Luo, Tse-Shun Huang, Bertrand Z Yeung, Efthymia Papalexi, et al. Scalable, multimodal profiling of chromatin accessibility, gene expression and protein levels in single cells. *Nature Biotechnology*, 39(10):1246–1258, 2021.

Vanessa M Peterson, Kelvin Xi Zhang, Namit Kumar, Jerelyn Wong, Lixia Li, Douglas C Wilson, Renee Moore, Terrill K McClanahan, Svetlana Sadekova, and Joel A Klappenbach. Multiplexed quantification of proteins and transcripts in single cells. *Nature Biotechnology*, 35(10):936–939, 2017.

Peter J Rousseeuw. Silhouettes: a graphical aid to the interpretation and validation of cluster analysis. *Journal of Computational and Applied Mathematics*, 20:53–65, 1987.

Alexander Strehl and Joydeep Ghosh. Cluster ensembles - a knowledge reuse framework for combining multiple partitions. *Journal of Machine Learning Research*, 3(Dec):583–617, 2002.

Jingshu Wang, Divyansh Agarwal, Mo Huang, Gang Hu, Zilu Zhou, Chengzhong Ye, and Nancy R Zhang. Data denoising with transfer learning in single-cell transcriptomics. *Nature Methods*, 16(9):875–878, 2019.

### Supplementary Figures

**A** DOGMA

| Method | Biological Variation Preservation (BVC) | | | | | Batch Effect Removal (BER) | | | | | Aggregate Score | | |
|---|---|---|---|---|---|---|---|---|---|---|---|---|---|
| | ILS | Kmeans NMI | Kmeans ARI | cASW | cList | bASW | iList | KBET | GC | PCR | BER | BVC | Overall |
| scProca | 0.52 | 0.60 | 0.58 | 0.52 | 0.97 | 0.95 | 0.85 | 0.75 | 0.81 | 0.99 | 0.87 | 0.64 | 0.73 |
| UNIMF | 0.50 | 0.55 | 0.42 | 0.53 | 0.98 | 0.88 | 0.75 | 0.73 | 0.75 | 0.89 | 0.80 | 0.59 | 0.68 |
| totalVI | 0.52 | 0.57 | 0.54 | 0.51 | 0.97 | 0.94 | 0.54 | 0.52 | 0.76 | 0.99 | 0.75 | 0.62 | 0.67 |
| StabMap | 0.51 | 0.60 | 0.52 | 0.51 | 0.96 | 0.90 | 0.70 | 0.59 | 0.57 | 1.00 | 0.75 | 0.62 | 0.67 |
| scArches | 0.53 | 0.58 | 0.55 | 0.52 | 0.98 | 0.93 | 0.44 | 0.49 | 0.79 | 0.98 | 0.73 | 0.63 | 0.67 |
| scMoMaT | 0.51 | 0.55 | 0.52 | 0.53 | 0.97 | 0.85 | 0.24 | 0.39 | 0.73 | 0.72 | 0.58 | 0.62 | 0.60 |

**B** REAP

| Method | Biological Variation Preservation (BVC) | | | | | Batch Effect Removal (BER) | | | | | Aggregate Score | | |
|---|---|---|---|---|---|---|---|---|---|---|---|---|---|
| | ILS | Kmeans NMI | Kmeans ARI | cASW | cList | bASW | iList | KBET | GC | PCR | BER | BVC | Overall |
| scProca | 0.58 | 0.79 | 0.83 | 0.53 | 0.99 | 0.95 | 0.88 | 0.90 | 0.83 | 0.99 | 0.93 | 0.82 | 0.82 |
| totalVI | 0.59 | 0.82 | 0.87 | 0.55 | 0.99 | 0.82 | 0.84 | 0.74 | 0.84 | 1.00 | 0.87 | 0.76 | 0.81 |
| scArches | 0.58 | 0.79 | 0.84 | 0.53 | 0.99 | 0.82 | 0.86 | 0.67 | 0.87 | 0.98 | 0.86 | 0.75 | 0.79 |
| UNIMF | 0.59 | 0.76 | 0.80 | 0.60 | 1.00 | 0.82 | 0.83 | 0.76 | 0.83 | 0.96 | 0.86 | 0.75 | 0.79 |
| StabMap | 0.62 | 0.68 | 0.59 | 0.63 | 1.00 | 0.82 | 0.55 | 0.59 | 0.81 | 1.00 | 0.77 | 0.71 | 0.73 |
| scMoMaT | 0.49 | 0.31 | 0.25 | 0.51 | 0.96 | 0.82 | 0.00 | 0.02 | 0.51 | 0.00 | 0.25 | 0.50 | 0.40 |

**C** SLN

| Method | Biological Variation Preservation (BVC) | | | | | Batch Effect Removal (BER) | | | | | Aggregate Score | | |
|---|---|---|---|---|---|---|---|---|---|---|---|---|---|
| | ILS | Kmeans NMI | Kmeans ARI | cASW | cList | bASW | iList | KBET | GC | PCR | BER | BVC | Overall |
| UNIMF | 0.52 | 0.59 | 0.41 | 0.52 | 0.99 | 0.96 | 0.91 | 0.90 | 0.73 | 0.77 | 0.85 | 0.61 | 0.71 |
| scProca | 0.55 | 0.64 | 0.43 | 0.53 | 0.99 | 0.97 | 0.90 | 0.90 | 0.87 | 0.36 | 0.80 | 0.63 | 0.70 |
| StabMap | 0.54 | 0.63 | 0.42 | 0.54 | 0.99 | 0.96 | 0.54 | 0.81 | 0.68 | 1.00 | 0.80 | 0.62 | 0.69 |
| scArches | 0.57 | 0.65 | 0.40 | 0.54 | 1.00 | 0.96 | 0.87 | 0.85 | 0.87 | 0.00 | 0.71 | 0.63 | 0.66 |
| totalVI | 0.56 | 0.65 | 0.40 | 0.54 | 1.00 | 0.96 | 0.86 | 0.82 | 0.88 | 0.00 | 0.70 | 0.63 | 0.66 |
| scMoMaT | 0.48 | 0.50 | 0.28 | 0.50 | 0.99 | 0.76 | 0.00 | 0.06 | 0.54 | 0.00 | 0.27 | 0.55 | 0.44 |

**D** BMMC

| Method | Biological Variation Preservation (BVC) | | | | | Batch Effect Removal (BER) | | | | | Aggregate Score | | |
|---|---|---|---|---|---|---|---|---|---|---|---|---|---|
| | ILS | Kmeans NMI | Kmeans ARI | cASW | cList | bASW | iList | KBET | GC | PCR | BER | BVC | Overall |
| UNIMF | 0.53 | 0.73 | 0.62 | 0.50 | 1.00 | 0.86 | 0.57 | 0.45 | 0.70 | 0.83 | 0.68 | 0.68 | 0.68 |
| scProca | 0.54 | 0.70 | 0.59 | 0.53 | 1.00 | 0.93 | 0.46 | 0.28 | 0.83 | 0.81 | 0.66 | 0.67 | 0.67 |
| StabMap | 0.54 | 0.73 | 0.64 | 0.58 | 1.00 | 0.86 | 0.06 | 0.18 | 0.74 | 1.00 | 0.57 | 0.70 | 0.65 |
| totalVI | 0.56 | 0.74 | 0.65 | 0.54 | 1.00 | 0.88 | 0.12 | 0.13 | 0.84 | 0.85 | 0.57 | 0.70 | 0.65 |
| scArches | 0.56 | 0.71 | 0.57 | 0.54 | 1.00 | 0.88 | 0.08 | 0.13 | 0.84 | 0.76 | 0.54 | 0.68 | 0.62 |
| scMoMaT | 0.54 | 0.51 | 0.33 | 0.51 | 0.99 | 0.76 | 0.00 | 0.00 | 0.70 | 0.00 | 0.30 | 0.58 | 0.47 |

**Supplementary Figure S1.** Comprehensive quantitative metrics of scProca and five other integration methods on the (A) DOGMA, (B) REAP, (C) SLN, and (D) BMMC datasets.

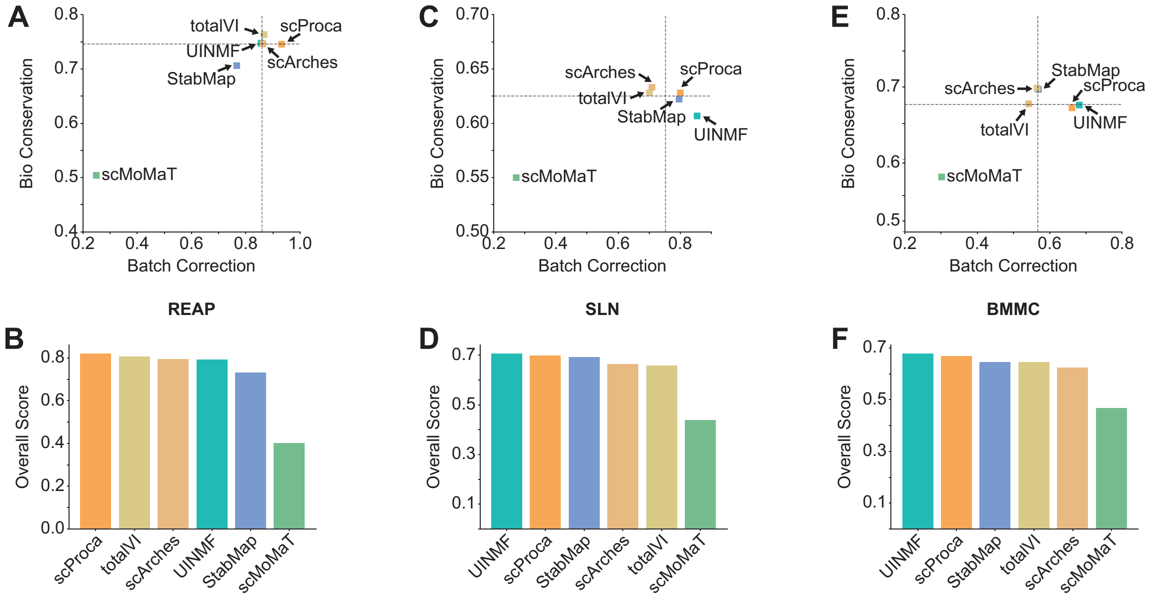


**Supplementary Figure S2.** (A) The scatter plot show biological variation conservation versus batch effect removal scores for integration, and (B) overall integration scores across all methods on the REAP dataset. (C) The scatter plot showing biological variation conservation versus batch effect removal scores for integration, and (D) overall integration scores across all methods on the SLN dataset. (E) The scatter plot showing biological variation conservation versus batch effect removal scores for integration, and (F) overall integration scores across all methods on the BMMC dataset. The dashed lines of scatter plots represent the median scores of all methods.

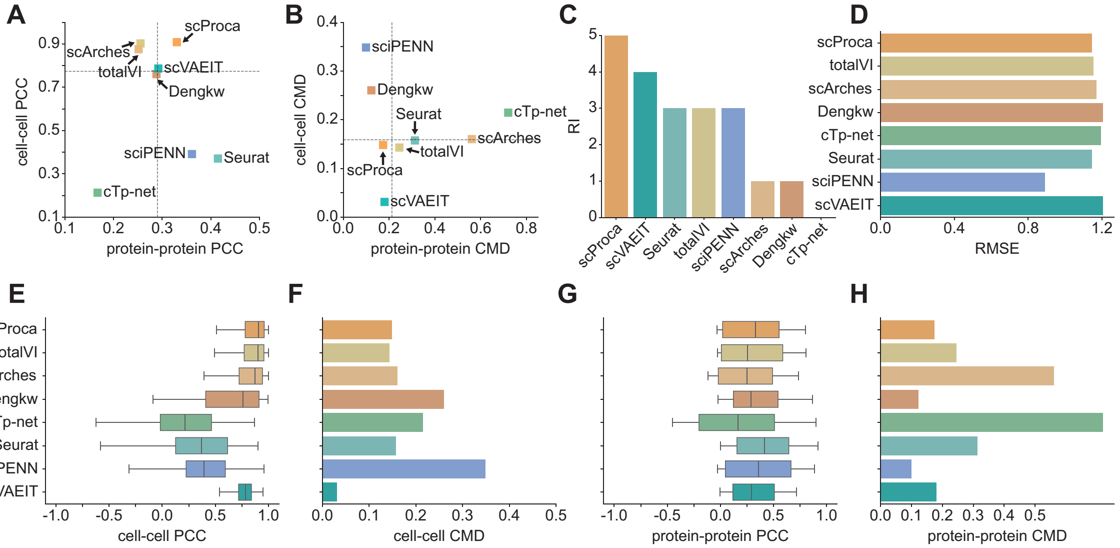


**Supplementary Figure S3:** Scatter plots showing (A) cell-cell PCC versus protein-protein PCC and (B) cell-cell CMD versus protein-protein CMD between the ADT measurements of CITE-seq cells and the predicted ADT measurements of scRNA-seq cells obtained by scProca and seven other generation, imputation, or prediction methods on the REAP dataset. The dashed lines are the median values of all methods. (C) Overall RI scores across all methods. (D) RMSE, (E) cell-cell PCC, (F) protein-protein PCC, (G) cell-cell CMD, and (H) protein-protein CMD values across all methods.

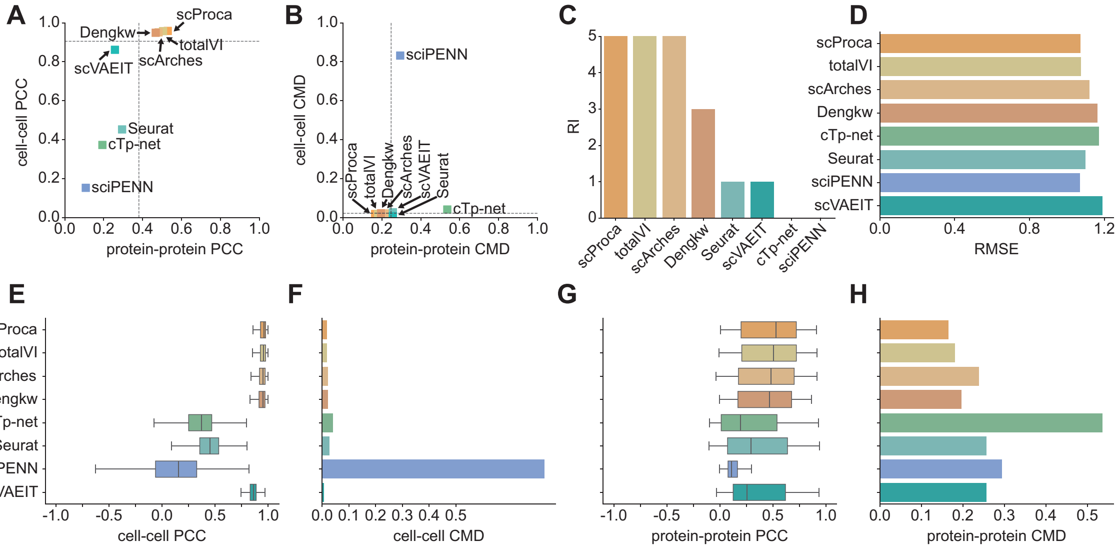


**Supplementary Figure S4:** Scatter plots showing (A) cell-cell PCC versus protein-protein PCC and (B) cell-cell CMD versus protein-protein CMD between the ADT measurements of CITE-seq cells and the predicted ADT measurements of scRNA-seq cells obtained by scProca and seven other generation, imputation, or prediction methods on the SLN dataset. The dashed lines are the median values of all methods. (C) Overall RI scores across all methods. (D) RMSE, (E) cell-cell PCC, (F) protein-protein PCC, (G) cell-cell CMD, and (H) protein-protein CMD values across all methods.

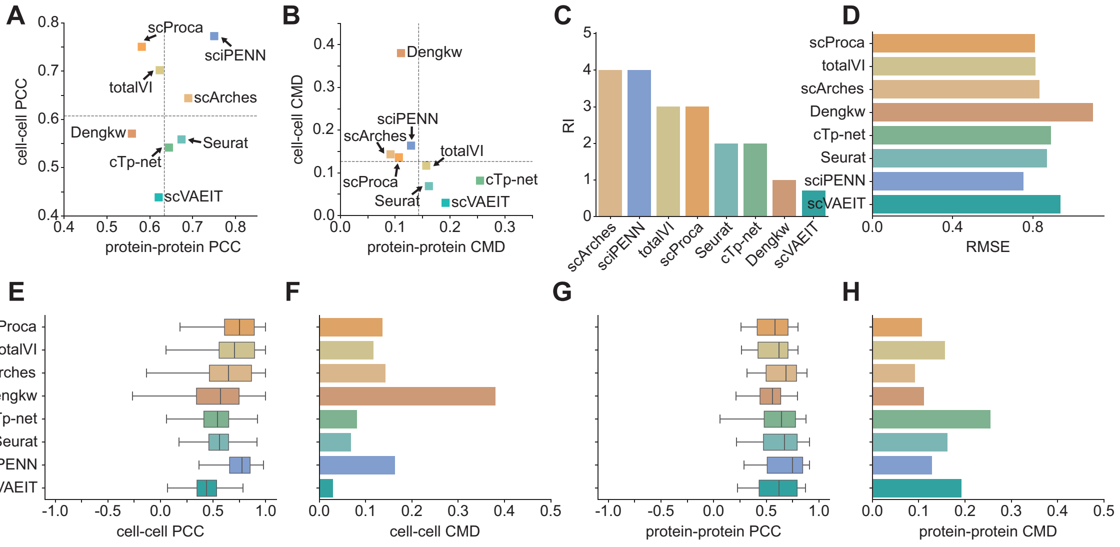


**Supplementary Figure S5:** Scatter plots showing (A) cell-cell PCC versus protein-protein PCC and (B) cell-cell CMD versus protein-protein CMD between the ADT measurements of CITE-seq cells and the predicted ADT measurements of scRNA-seq cells obtained by scProca and seven other generation, imputation, or prediction methods on the BMMC dataset. The dashed lines are the median values of all methods. (C) Overall RI scores across all methods. (D) RMSE, (E) cell-cell PCC, (F) protein-protein PCC, (G) cell-cell CMD, and (H) protein-protein CMD values across all methods.

**A**

| | Biological Variation Preservation (BVC) | | | | | Batch Effect Removal (BER) | | | | | Aggregate Score | | |
|---|---|---|---|---|---|---|---|---|---|---|---|---|---|
| **Scenario** | ILS | Kmeans NMI | Kmeans ARI | cASW | cList | bASW | iList | KBET | GC | PCR | BER | BVC | Overall |
| 1 v.s. 3 | 0.55 | 0.69 | 0.56 | 0.53 | 0.99 | 0.97 | 0.59 | 0.89 | 0.85 | 0.99 | 0.86 | 0.66 | 0.74 |
| 2 v.s. 2 | 0.55 | 0.64 | 0.43 | 0.53 | 0.99 | 0.97 | 0.90 | 0.90 | 0.87 | 0.36 | 0.80 | 0.63 | 0.70 |

**B**

| | Generation performance | | | | |
|---|---|---|---|---|---|
| **Scenario** | RMSE | cell-cell PCC | cell-cell CMD | protein-protein PCC | protein-protein CMD |
| 1 v.s. 3 | 1.04 | 0.95 | 0.02 | 0.48 | 0.17 |
| 2 v.s. 2 | 1.07 | 0.96 | 0.02 | 0.53 | 0.16 |

**Supplementary Figure S6:** Comprehensive quantitative metrics of scProca on the SLN dataset under two scenarios: '1 v.s. 3' refers to the case where only SLN111-D1 serves as CITE-seq data, while the others serve as scRNA-seq data and '2 v.s. 2' refers to the case where both SLN111-D1 and SLN111-D2 serve as CITE-seq data, while both SLN208-D1 and SLN208-D2 serve as scRNA-seq data. (A) Integration metrics. (B) Generation metrics.
